# Global phylogenetic analysis of *Escherichia coli* and plasmids carrying the *mcr-1* gene indicates bacterial diversity but plasmid restriction

**DOI:** 10.1101/136804

**Authors:** Sébastien Matamoros, Jarne M. van Hattem, Maris S. Arcilla, Niels Willemse, Damian C. Melles, John Penders, Trung Nguyen Vinh, Ngo Thi Hoa, Menno D. de Jong, Constance Schultsz, Martin C.J. Bootsma, Perry J. van Genderen, Abraham Goorhuis, Martin P. Grobusch, Nicky Molhoek, Astrid M.L. Oude Lashof, Ellen E. Stobberingh, Henri A. Verbrugh

## Abstract

To understand the dynamics behind the worldwide spread of the *mcr-1* gene, we determined the population structure of *Escherichia coli* and of mobile genetic elements (MGEs) carrying the *mcr-1* gene. After a systematic review of the literature we included 65 *E. coli* whole genome sequences (WGS), adding 6 recently sequenced travel related isolates, and 312 MLST profiles. We included 219 MGEs described in 7 Enterobacteriaceae species isolated from human, animal and environmental samples. Despite a high overall diversity, 2 lineages were observed in the *E. coli* population that may function as reservoirs of the *mcr-1* gene, the largest of which was linked to ST10, a sequence type known for its ubiquity in human faecal samples and in food samples. No genotypic clustering by geographical origin or isolation source was observed. Amongst a total of 13 plasmid incompatibility types, the Incl2, lncX4 and IncHI2 plasmids accounted for more than 90% of MGEs carrying the *mcr-1* gene. We observed significant geographical clustering with regional spread of lncHI2 plasmids in Europe and Incl2 in Asia. These findings point towards promiscuous spread of the *mcr-1* gene by efficient horizontal gene transfer dominated by a limited number of plasmid incompatibility types.

## Introduction

Antimicrobial resistance (AMR) represents a growing threat to global health ^1^. With barely any new antimicrobial drugs in development^2^, limiting the spread of AMR is key in order to maintain current treatment options ^3^.

Colistin is an antibiotic of the polymyxin class, discovered in 1950 and effective against Gram-negative bacteria ^4^. The emergence of multidrug-resistant Gram-negative bacteria, especially those producing carbapenemases, has reintroduced colistin as a last resort antibiotic for the treatment of severe infections^5^. In contrast to its limited use in humans, colistin is widely used in food-producing animals ^6^. While colistin resistance was long thought to be caused by chromosomal mutations only ^7^, the emergence of plasmid-mediated resistance, conferred by the mobilized colistin resistance *(mcr-1)* gene, was recently reported ^8^. This gene encodes for a protein of the phosphoethanolamine transferase enzyme family, and its expression results in the addition of a phosphoethanolamine to lipid A, the target of colistin, decreasing the interaction between colistin and the bacterial lipopolysaccharide^8^. Since its discovery in 2015 in China, this gene has been described in several bacterial species that were isolated from animals, animal food products, humans and environmental samples from around the world ^9–13^. Our previous study in travellers indicated acquisition of *mcr-1* carrying bacteria by healthy individuals during travel to destinations around the world, potentially related to food exposure, as well as rapid clearance after return ^14^. It has been suggested that *mcr-1* has spread from food animals to humans ^8,15–17^, but there is a lack of comparison of *mcr-1* carrying isolates on a global level to support this hypothesis.

We studied the global population structure as well as the geographic and host distribution of *mcr-1-* carrying *Escherichia coli,* and mobile genetic elements (MGEs), to establish the population structure and to assess whether the spread of the *mcr-1* gene is linked to clonal dissemination or transmission of MGEs from animal, human, or environmental sources within geographic regions.

## Results

### Literature search

A systematic review of the literature on *mcr-1,* published until 1 January 2017 resulted in the inclusion of 95 articles, representing a total of 410 entries (whole genome sequences, MLST profiles, and/or plasmid types) for analysis (See detailed methods and results in Supplementary data, Supplementary Figure 1 and Supplementary Table 1).

### Population structure

#### Whole genome sequencing (WGS)

The genomes of 65 *mcr-1*-carrying *E. coli* were analysed, including 6 genomes obtained from *E. coli* isolated from travellers that were sequenced for the purpose of the present study. Isolates originated from Asia (n=36; 55.4%), Europe (n=20; 30.8%), North-America (n=4; 6.2%), South-America (n=4; 6.2%) and Africa (n=1; 1.5%). 45 were of animal origin (69.2%), 19 of human origin (29.2%) and one strain (1.5%) was isolated from water (Supplementary Table 1).

The average size of the genomes (all contigs in each assembly, representing chromosomes and plasmids) of these 65 isolates was 4.9 Mbp, with a median number of genes identified of 4785 (ranging from 4266 to 7083), representing a pangenome of 23248 genes and a core genome (defined by genes present in at least 99% of the isolates) of 2216 genes. An unbiased analysis of the population structure was performed using a Bayesian approach with the BAPS software ^18^, based on the nucleotide alignment of the core genome sequences. It revealed the presence of 5 distinct phylogenetic clusters (Figure 1; Supplementary Figure 2; Supplementary Table 1). The largest cluster (cluster 1) consisted of 26 isolates from 16 different STs (26/65; 40.0%) and the second cluster of 24 isolates from 15 different STs (36.9%). No significant relationship between clustering (BAPS) and geographical origin or isolation source was observed (χ^2^-test) (Figure 1) except that all 5 isolates that belong to BAPS cluster 3 are from Europe. Twenty isolates showed less than 10 SNPs/Mbp difference and were considered clonally related (Supplementary Tables 2 and 3).

**Figure 1:**
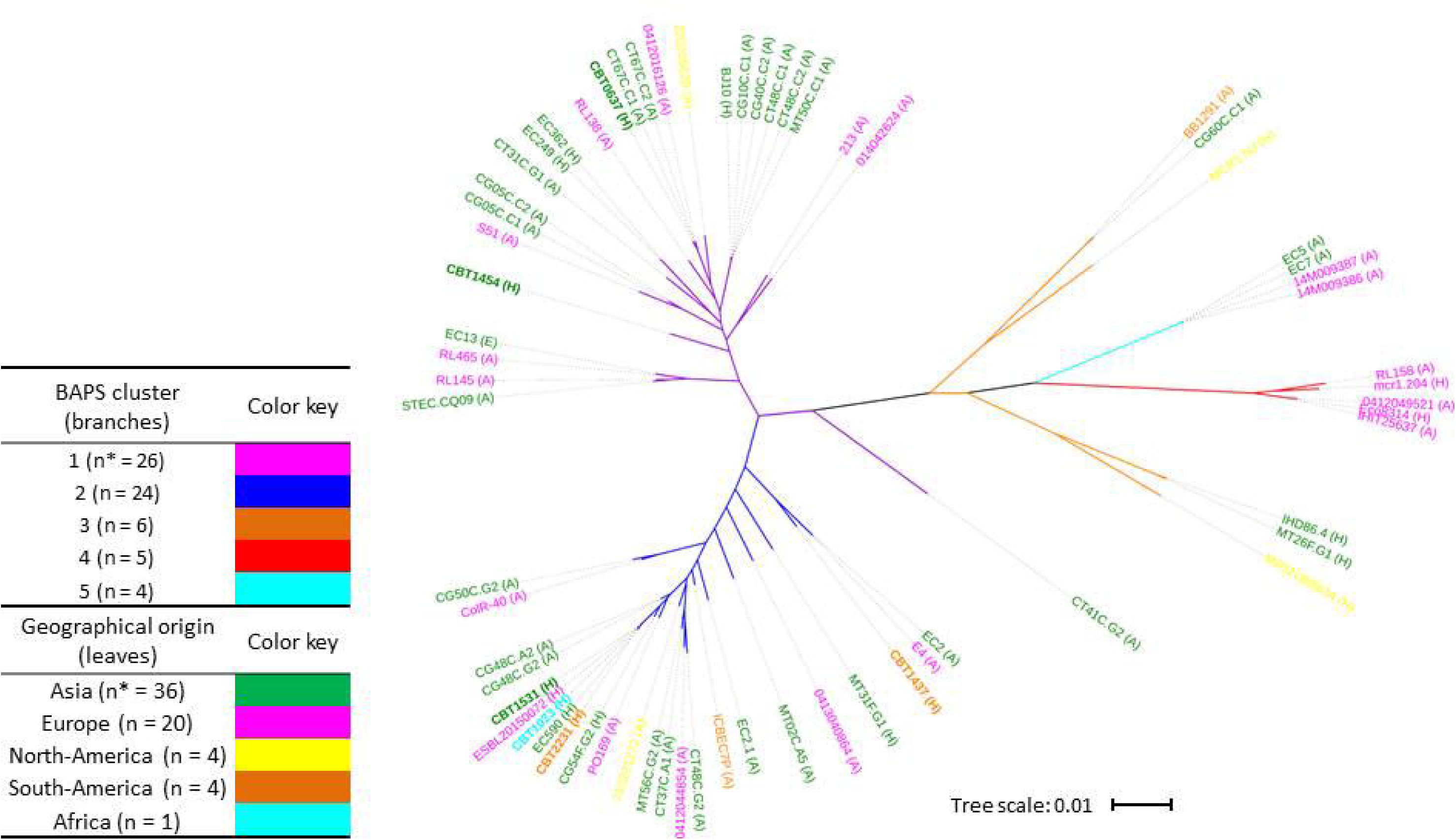
Maximum-likelihood tree based on concatenated core genome sequences of 65 *mcr-1-* carrying £ *coli* isolates. Branch colours indicate phylogenetic clusters as determined by BAPS. Isolates from ST10, ST165 and closely related isolates are all grouped in the BAPS cluster 2 (dark blue). Leaf colours (isolates identifiers) indicate geographical region of origin. Isolation source is indicated in brackets: A = animal or meat; H = human; E = environment. The 6 travellers’ isolates that were sequenced for this study are highlighted in bold and names start with CBT. Tree scale in number of substitutions per site. *number of isolates.

#### Multilocus sequence typing (MLST)

For 312 *E. coli* isolates originating from 69 studies, a MLST profile was published or could be deduced from the corresponding WGS. Of these, 206 were isolated from animals or animal products (66.0%), 101 were isolated from humans (32.4%), including the 6 travel acquired isolates, and 5 from the environment (1.6%). 141 Isolates from 25 studies (141/312; 45.2%) originated from Asia and 125 isolates from 25 studies (40.1%) from Europe together accounting for 85.3% of all included isolates. The isolates represented 112 unique sequence types (STs) with ST10 being most common, comprising 40/312 (12.8%) isolates originating from Africa, Asia, Europe and South-America. eBURST analysis ^19^ was performed on all isolates included in the study, to identify their genetic relatedness based on their MLST profiles. Three main clusters were identified, for which the predicted founders, i.e. the ST in a cluster from which all other SLVs and DLVs in the cluster have most likely diversified ^19^, were ST10, ST1114 and ST410. The largest cluster contained all 40 ST10 isolates and an additional 46 isolates in 21 STs that were single (SLV) or double locus variants (DLV) of ST10 (86/312; 27.6%) (Supplementary Figure 3). The predicted founder of the second largest cluster was ST1114, a SLV of ST165 and ST100, and included 19 isolates belonging to 7 different STs (5.4%), while the third cluster was centred on ST410 and included 14 isolates from 3 different STs (4.5%).

A maximum-likelihood tree based on concatenated MLST gene sequences showed a main clade of 128 isolates (represented by blue branches in Figure 2 and Supplementary Figure 4; bootstrap value of the main branch = 0.98), including most, but not all, isolates from the eBURST clusters of ST10 and ST1114 (Supplementary Figure 4A). All isolates from these 2 eBURST clusters for which a WGS was available were grouped in BAPS cluster 2. Similarly, all the isolates from the eBURST cluster ST410 grouped into BAPS cluster 1, along with 6 isolates from ST155. Seven isolates belonged to the globally successful extra-intestinal pathogenic *E. coli* clone ST131 (Supplementary Figure 4A). As observed in the WGS analysis, animal isolates were interspersed with isolates from humans and the environment throughout the tree, as were isolates from different continents indicating a lack of clustering by isolation source or geographical origin (Figure 2). Similarly, no clustering by health status of the host was observed (Supplementary Figure 4B).

**Figure 2:**
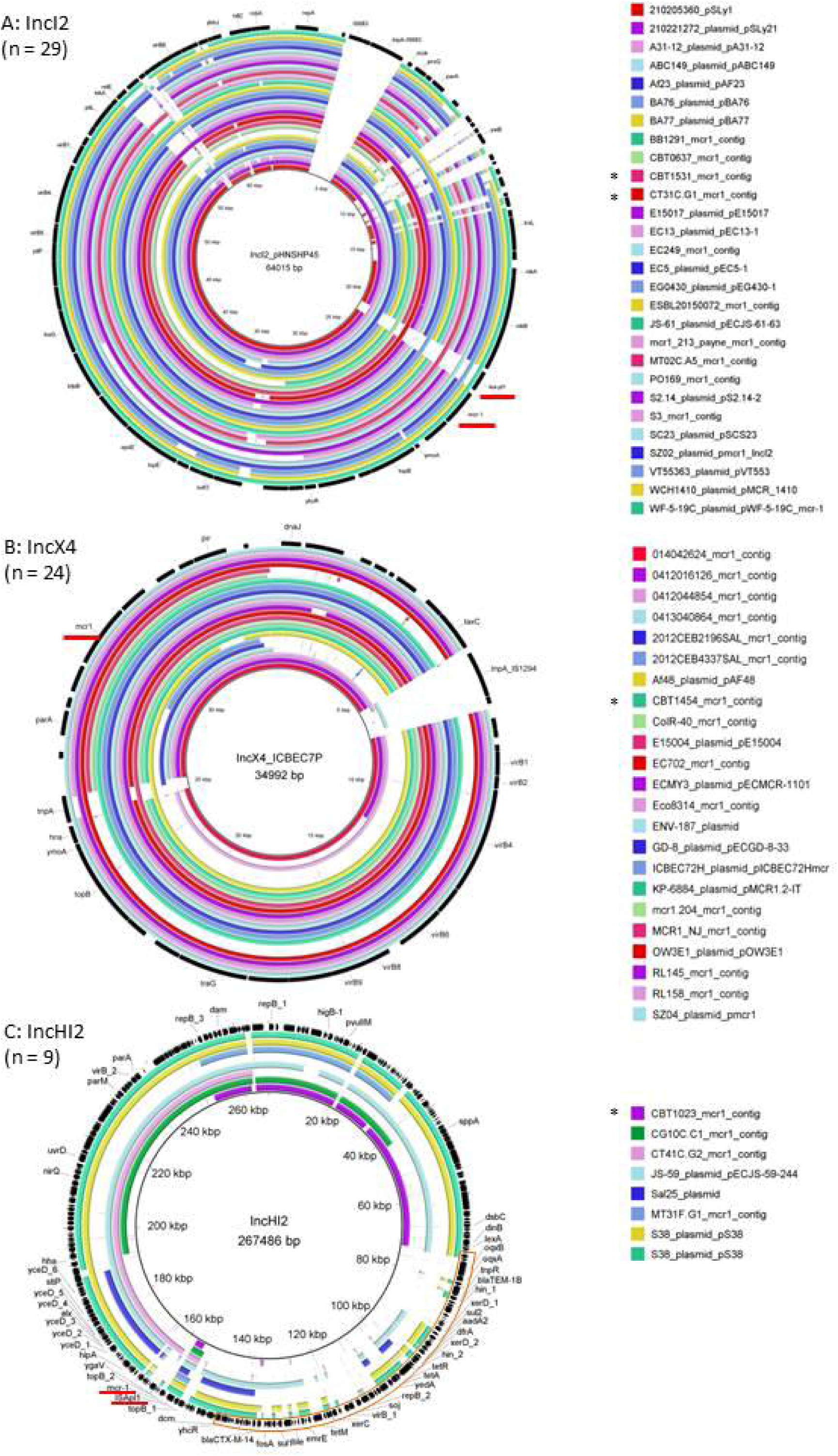
Phylogeny of the *mcr-1-*carrying *E. coll* isolates. Maximum-likelihood tree based on concatenated MLST gene sequences, mid-point rooted. Inner coloured circle: isolation source; outer circle: region of origin. Stars indicate the isolates from which a whole genome sequence was available. The 6 travellers’ isolates that were sequenced for this study are highlighted in green. Bootstrap values between 0.9 and 1 are indicated by red triangles (size proportional to bootstrap value). The blue branches represent the main clade of 128 isolates including most isolates from STIO.Tree scale in number of substitutions per site. See Supplementary Figure 4A for additional information on the relationship between STs, eBURST clustering and WGS BAPS clustering.

### Mobile genetic elements

The plasmid incompatibility group of the *mcr-1-*carrying plasmids could be determined for 217 Enterobacteriaceae isolates from 7 different species (*Escherichia sp., Salmonella* sp., *Klebsiella* sp., *Cronobacter* sp., *Enterobacter* sp., *Kluyvera* sp. and *Shigella* sp.), representing a total of 219 plasmids since 2 isolates carried 2 different plasmids carrying the *mcr-1* gene (Table 1). These plasmids were described in 71 studies (1 to 33 plasmids per study, average = 3.1). In addition, the gene was integrated in the chromosome of 6 isolates. The incompatibility group could not be determined for 27 of the 65 isolates for which WGS was available. Similarly the plasmid type was not available for 182 of the 312 isolates included in the MLST analysis. A total of 14 different plasmid incompatibility groups were identified. 198/219 (90.4%) of the identified plasmids belonged to one of 3 incompatibility groups: IncX4 (77/219 plasmids, 35.2%), IncI2 (76/219 plasmids, 34.7%) and IncHI2 (45/219 plasmids, 20.5%). 50/76 Incl2 plasmids (65.8%) originated from Asia and 33/45 IncHI2 plasmids (73.3%) from Europe. IncX4 plasmids were more evenly distributed: 44/77 (57.1%) were recovered from Europe, 29 from Asia (37.7%) and 4 from other regions (5.2%). Observed proportions were significantly different from expected for Incl2 (χ^2^-test, p < 0.001) and IncHI2 (p < 0.001) but not for IncX4. The distribution of these 3 plasmid types was not significantly different from expected between animal (χ^2^-test, p = 0.24), human (p = 0.88) and environmental sources (p = 0.38). Isolates from the BAPS groups 1 and 2 carried plasmids from the 3 major types in similar proportions (Supplementary Figure 5A; Supplementary Table 1). Isolates from the eBURST clusters of ST10 carried plasmids belonging to 7 different incompatibility groups, including IncHI2, IncI2 and IncX4. No clustering of plasmid type with MLST phylogeny was observed either (Supplementary Figure 5B; Supplementary Table 1).

**Table 1:**
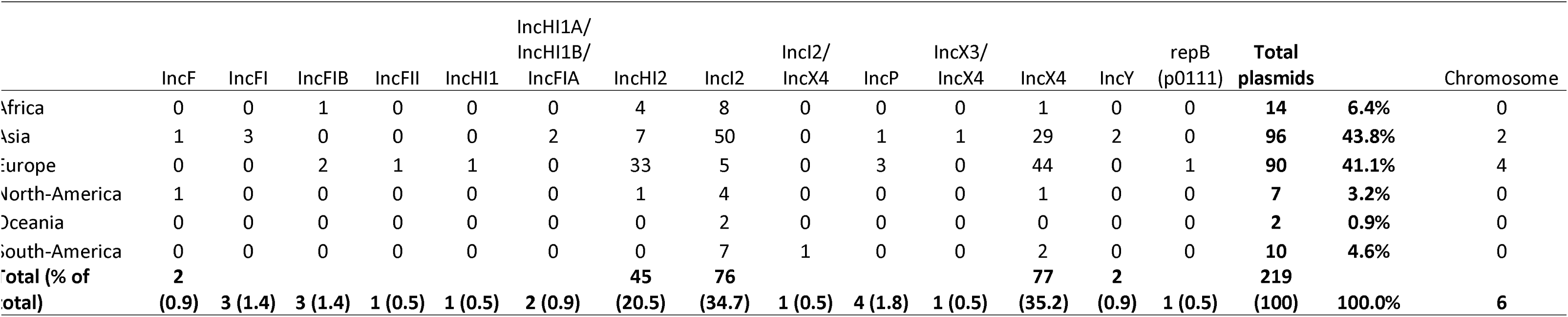
Incompatibility types of *mcr-1* carrying plasmids and distribution by geographical regions

Figure 3 shows the alignment of the complete sequences or contigs from Incl2 (panel A), IncX4 (B) and lncHI2 (C) incompatibility group plasmids. IncHI2 plasmids had the largest size, with sequence lengths up to 267486 bp.

**Figure 3:**
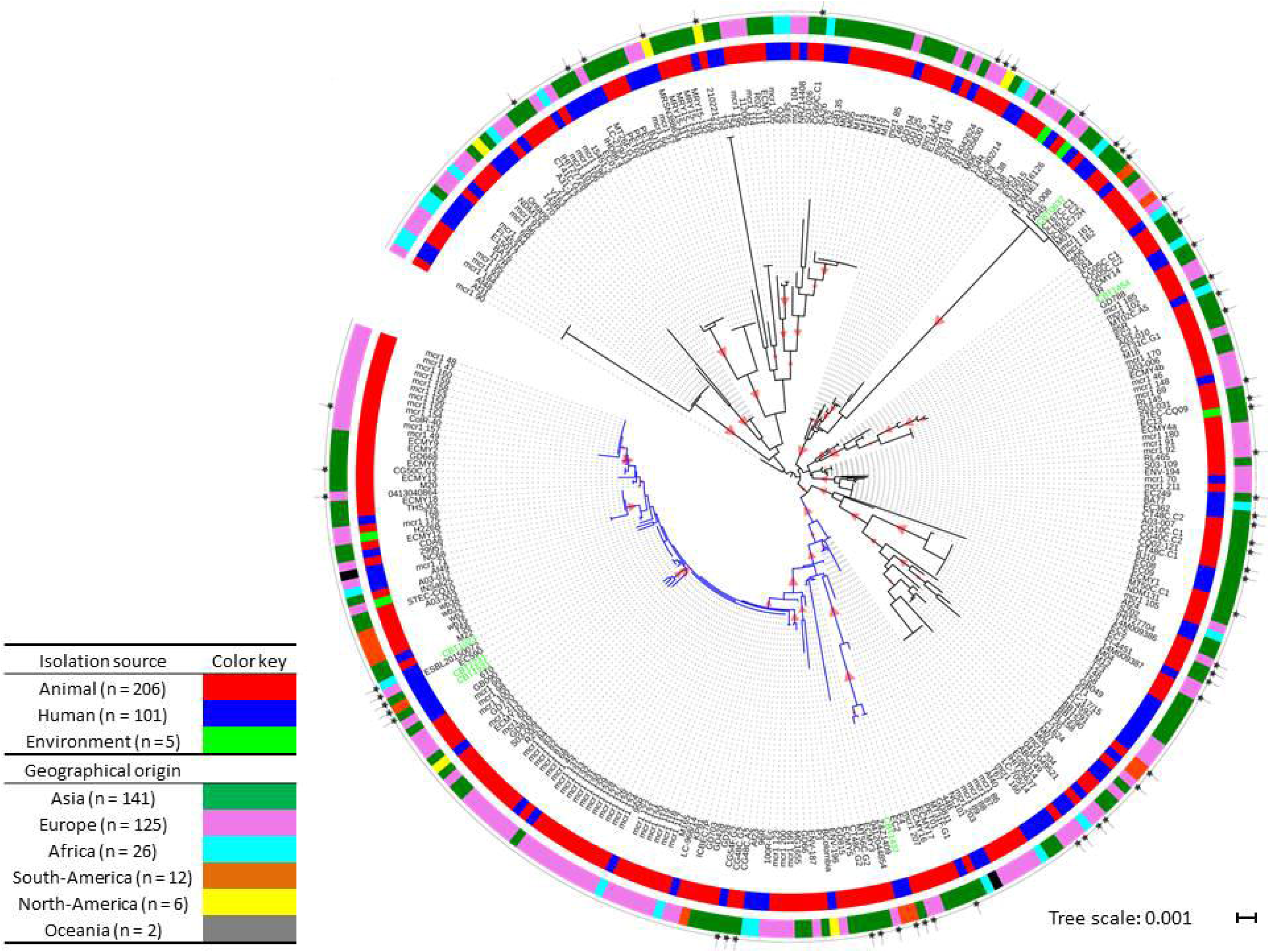
Alignment of mcr-T-containing plasmids and contigs. Panel A: IncI2 plasmids (n = 29); panel B: IncX4 (n = 24); panel C: IncHI2 (n = 9). Black outer ring: plasmid used as reference for the alignment; name and size of the reference indicated in the middle of each panel. Plasmid names followed by "_mcrl_contig" refer to assembled contigs from whole genome sequences. Other names refer to plasmid sequences deposited in online databases. The *mcr-1* gene and ISA-pIl location are underlined in red. Plasmids indicated with an asterisk are from the 6 travellers’ isolates that were sequenced for this study. Panel C: Putative MDR cassette is highlighted in orange.

The ISA*p/2* transposon element situated upstream of the *mcr-1* gene was present in 7/9 (77.8%) IncHI2 plasmids, but only in 11/29 (37.9%) of IncI2 plasmids and completely missing in all of the 24 reported IncX4 plasmids (Figure 3). In the isolates from travellers the ISA*p/l* transposon was identified in 3 out of our 6 *mcr-1*-carrying contigs including one isolate from a traveler to Asia (ST101, IncI2 incompatibility group), one to Africa (ST744, IncHI2) and one to South-America (ST744, incompatibility group not identified).

### Antimicrobial resistance genes

All sequences of the *mcr-1* gene collected in the present study were 100% identical to the original sequence described by Liu *et al*.^8^.

Multiple resistance genes were detected in most of the studied isolates (Supplementary Results). The florfenicol resistance gene *floR* was present in 32 (49.2%) isolates; in 22 of 45 isolates from animals (48.9%) and 10 of 19 isolates from humans (52.6%). The *baeR* and *baeS* genes, encoding novobiocin resistance, were found in 64 (98.5%) and 65 (100%) isolates, respectively.

Plasmid analysis from the WGS data showed that 4 of the 29 *mcr-1*-carrying IncI2 plasmids (13.8%) contained an additional ESBL gene. IncHI2 plasmids (n=9) carried between 0 and 12 additional AMR genes. In particular, 4 plasmids carried CTX-M ESBL genes and 2 carried the *floR* gene. In 4 out of the 9 IncHI2 plasmids analysed in this study, the *mcr-1* gene was shown to be integrated alongside a large multi-drug resistance (MDR) gene cassette (Figure 3C). None of the IncX4 plasmids carried additional AMR genes.

## Discussion

Analysis of all reported WGS of *mcr-1*-carrying isolates shows that the population of *E. coli* is highly diverse, but is dominated by 2 large groups of related isolates. Most of the isolates from BAPS group 2 grouped into a MLST cluster centred on ST10. An overrepresentation of isolates related to ST10 and ST165 (a SLV of ST1114) in *mcr-1-*carrying *E. coli* isolates was previously reported at a smaller scale in isolates from European farm animals ^20^. *E. coli* ST10 and closely related STs are frequently recovered from food and human intestinal samples and studies have shown a higher prevalence of plasmid-carried AMR genes in ST10, including CTX-M ESBL genes, compared to other STs ^21–24^. The second BAPS group of interest in our study included isolates belonging to ST155. This ST has been described as a major vector of spread of ESBL genes from animals to humans ^25^. It is thus possible that zoonotic transmission lead to the spread of the *mcr-1* gene, as has been suggested in studies from China and Vietnam ^16,17^, notably through the 2 main phylogenetic clusters identified in this study.

Additionally, we found clonally related isolates, including some belonging to ST744, a SLV of ST10, carrying the *mcr-1* gene on different plasmid backbones and recovered from different continents (see Supplementary Results). These results point towards a worldwide dissemination *of mcr-1* driven mainly by highly promiscuous plasmids rather than the worldwide spread of one or more *mcr-1* carrying clones. We hypothesize that several populations of *E. coli* isolates, notably those related to ST10 or ST155, acquired the *mcr-1* gene due to their intrinsic ability of acquiring AMR genes and their high prevalence in humans and food animals. These populations of commensal isolates then may play a crucial role as a reservoir for this gene, which can explain their over-representation in the present study.

In the timeframe of our literature search, 3 *E. coli* strains carrying the *mcr-2* gene were isolated from animals in Belgium. These isolates belonged to ST10 (2 isolates) and ST167 which is a SLV of ST10 and carried the gene on an IncX4-type plasmid. No WGS data was available from these isolates ^26^. More data about the *mcr-2* gene is needed to assess its spread and determine if *E. coli* ST10 plays a similar role in its dissemination as it does for *mcr-1.*

More than 90% of published plasmid types carrying *mcr-1* genes belonged to either IncI2, IncX4 or IncHI2. Almost 75% of the isolates carrying an IncHI2 plasmid originated from Europe: 26 from animals and 7 from humans (Table 1 and Supplementary Table 1). In a traveller’s isolate acquired in Tunisia, the *mcr-1*-carrying plasmid was identified as an IncHI2-type backbone of the ST4 pMLST subtype which co-carried a CTX-M-1 ESBL gene (Figure 3C). This traveller reported consumption of beef, chicken and eggs during travel to Tunisia which can potentially be the source for the acquisition of the *mcr-1* positive isolate. When investigating the presence of the *mcr-1* gene in cephalosporin resistant *E. coli* isolates from chicken farms in Tunisia, Grami *et al.*^27^ found that all 37 *mcr-1*-carrying plasmids also belonged to the IncHI2-type, ST4 subtype and harboured CTX-M-1 genes. PFGE typing of the isolates harbouring this plasmid showed various bacterial genetic backgrounds. Interestingly, these chickens were all imported from France, either as adults or chicks. Other studies showed the presence of this IncHI2, CTX-M-1 and *mcr-1* combination in *Salmonella enterica* Typhimurium isolates from meat samples in Portugal from 2011 ^28,29^ and diarrhoeic veal calves in France ^30^. The IncHI2 subtype ST4 was also detected in an *E. coli* isolate from retail chicken breast in Germany ^31^ and the faecal sample of a veal calve from the Netherlands ^32^, suggesting widespread dissemination of this particular plasmid in European farm animals and possible transmission to humans.

The high prevalence of novobiocin *baeR* and *baeS* and florfenicol *floR* resistance genes ^33,34^ in the genomes of isolates of human and animal origin together with the fact that florfenicol and novobiocin are used almost exclusively in veterinary medicine further supports the potential role of food animals as an important reservoir of *mcr-1* containing bacteria and MGEs ^15^.

In contrast with the IncHI2 plasmids, 65.8% of all IncI2 plasmids recovered so far originated from Asia, with a much lower prevalence in *mcr-1* carrying Enterobacteriaceae from other regions. Taken together, these elements point toward a more regional circulation and dissemination of the *mcr-1-*carrying plasmids IncI2 and IncHI2.

We found the ISA*p/l* transposon element associated with the *mcr-1* gene, as originally described by Liu *et al.*^8^ to be present in a minority of studied plasmids and contigs. However, since some of the *mcr-1-*carrying contigs were obtained by assembly of Illumina short reads from WGS data, we cannot exclude that some of these gaps are explained by an incomplete assembly of (plasmid) sequences. The ISA*p/1* transposon element is considered to be the main driver of horizontal gene transfer of the *mcr-1* gene and has been shown to be highly unstable in IncI2 plasmids ^35–37^. The absence of the ISA*pl1* transposon element in *mcr-1*-carrying IncX4 plasmids as described here has recently been proposed to be essential for the maintenance of the *mcr-1* gene in this particular backbone, but the exact mechanism still requires further investigation ^38^.

WGS analysis provided in-depth information about the *mcr-1*-carrying *E. coli* isolates and their phylogenetic relationship, but the number of available genomes was limited. On the other hand, whilst MLST data have a lower resolution, the higher number of available profiles allowed analysis of the isolates’ origin (geographical, source of isolation, diseased status of the host, etc.).

A limitation of our study is the potential for bias. The overrepresentation of isolates originating from Asia and Europe could be explained by a higher prevalence of *mcr-1* genes on these continents, but the effect of publication bias cannot be excluded. Isolates from North-America only represented 2.2% of the collection. Noteworthy, colistin, except for ophthalmic ointment, has never been marketed for use in animals in the United States ^39,40^.

Sampling bias should also be considered when several isolates with an identical ST are presented from a single study, as is the case for ST100 and ST752. Additionally, in the absence of a control population of mcr-1-negative isolates obtained from similar sources as the *mcr-1* positive isolates, results of analysis of population structures should be interpreted with caution. Because many studies screened existing collections of (resistant) isolates for colistin resistance or presence of *mcr-1,* selection bias has probably been introduced.

The findings in this study suggests that the *mcr-1* gene has locally and globally disseminated through MGEs that are mainly IncHI2, IncI2 and IncX4 plasmids and provides additional support for the hypothesis of the animal reservoir, that is driven by the use of colistin in livestock, as a source of *mcr-1* in humans. A global ban of colistin use in animals to preserve colistin for use in human medicine seems therefore justified.

## Material and Methods

### Selection of isolates for whole genome sequencing

We subjected 6 *mcr-1* positive isolates that were collected as part of a prospective study (COMBAT study) aimed at studying acquisition of extended-spectrum β-lactamase (ESBL)-producing Enterobacteriaceae during travel to whole-genome sequencing ^14,41^. Additionally, we included 22 whole genome sequences of isolates from Vietnamese chickens and humans that were still unpublished when performing our literature search ^17^.

### Literature search

Relevant papers that published on *mcr-1* and *mcr-2* were identified in PubMed, Web of Science, Scopus, ScienceDirect and Google Scholar using the query 'mcr-1 OR mcr1 OR mcr-2 OR mcr2 OR (mcr AND colistin)’ (see Supplementary Material for full search strategies).To be able to study the associations between phylogeny, geographic distribution and isolation source we only included sequences from papers that provided sufficient metadata. As a consequence, plasmids and sequences that were deposited in online databases without metadata were not included in the analysis.

### Whole genome sequencing of *mcr-1*-positive *E. coli* isolates

Bacterial DNA was extracted from fresh pure cultures using the Qiagen DNeasy Blood and Tissue kit (Qiagen, Hilden, Germany). Library preparation was done according to manufacturer’s instruction (Illumina, San Diego, CA, USA) and sequenced using Illumina MiSeq technology with 150nt paired-end settings. Sequences have been deposited in the European Nucleotide Archive under the accession numbers SAME104030441 to SAME104030446.

### Bio-informatic analysis

#### MLST analysis

For each *mcr-1*-carrying *E. coli* isolate for which the ST or the whole genome sequence was available, the sequences of the corresponding alleles were downloaded from the *E. coli* MLST genes repository of the University of Warwick (http://mlst.warwick.ac.uk/mlst/dbs/Ecoli/handlers/getFileData/home/cbailster/mlst/zope/Extensions/gadfly/Ecoli/DB/) and concatenated. When STs of isolates were not described in literature, the ST was determined from available whole genomes using the online service provided by the Center for Genomic Epidemiology (https://cge.cbs.dtu.dk/services/MLST/) according to the Achtman MLST scheme ^42,43^. MLST clusters (STs and their single locus or double locus variants) were defined using e-burst V3 (http://eburst.mlst.net/v3/enterdata/single/)^19^ and goeBURST v1.2.1 ^44^ using only profiles from this study.

#### WGS and plasmid analysis

Raw sequence reads in fastq format or pre-assembled sequences in fasta format were downloaded from online databases for all available isolates (Supplementary Table 1). Additional sequences not yet deposited in online databases were requested from their respective authors. The quality of the raw sequence reads was checked using fastqc (http://www.bioinformatics.babraham.ac.uk/proiects/fastqc/), quasi^45^ and KmerFinder 2.0 (https://cge.cbs.dtu.dk/services/KmerFinder/) (see Supplementary Methods for more details). Reads were trimmed using Trimmomatic V0.33 ^46^. *De-novo* genome assembly was performed with SPAdes 3.9 ^47^ for Illumina short reads and with Canu v1.3 for PacBio long reads ^48^. Contigs of less than 500 bp long were removed from the genomes to improve the overall quality of the assembly. Size of the genomes was calculated by adding the length of all remaining contigs. Identification of open reading frames (ORFs) and gene contents in the assembled genomes *(de-novo* assemblies and pre-assembled sequences) was performed using Prokka v1.11 ^49^. Core genome analysis was performed with Roary v3.6.8 ^50^. Clustering of isolates was performed using the hierBAPS module of the Bayesian Analysis of Population Structure (BAPS) software v6.0 ^18^. The core genome alignment output provided by Roary was used as input for BAPS with 2 levels of hierarchy and a maximum number of cluster (K) of 10. The estimated number of clusters was 5 for both levels of hierarchy.

Sequences (concatenated MLST loci or concatenated core genes) were aligned using mafft v6.864b ^51^. The resulting alignment was used as input for calculation of distances and tree building using RAxML v8.1.6 ^52^. TMLST and WGS trees were visualized using iTOL v3.3.1 (http://itol.embl.de/)^53^.

Identification of plasmid incompatibility group and typing of IncHI2 plasmids were performed on assembled sequences *(de-novo* or pre-assembled) via the CGE online services PlasmidFinder v1.3 (https://cge.cbs.dtu.dk/services/PlasmidFinder/) and pMLST v1.4 (https://cge.cbs.dtu.dk/services/plVILST/)^54^. Alignment and visualization of plasmids was performed with BRIG v0.95 ^55^. The majority of the isolates and plasmids described in this study were sequenced using a short read technology (Illumina). This technology does not allow for a high quality assembly of the plasmids due to the high number of repeat regions present in these MGEs. Therefore no phylogenetic analysis of the *mcr-1*-carrying plasmids was conducted in this study.

Two different databases were used for identification of other antibiotic resistance genes: ResFinder (https://cge.cbs.dtu.dk/services/ResFinder^56^) was used to detect acquired resistance genes commonly located on mobile genetic elements (MGEs) and CARD Resistance Gene Identifier (https://card.mcmaster.ca/analyze/rgi^57^) was used to detect chromosomal genes.

#### Statistics

The distribution of isolates and plasmids from different geographical origins and isolation sources was determined by a χ^2^-test comparing the expected distribution (proportions of the total studied population) to the observed proportions using GraphPad Prism6 (La Jolla, CA, USA).

## Acknowledgments

The authors would like to thank the team of the Department of Genome analysis of the AMC (in particular Dr. F. Baas and L. Koster) for their support in the sequencing bacterial genomes. Financial support for this study was provided by the European COMPARE project (http://www.compare-europe.eu/) under the European Union’s Horizon 2020 research and innovation programme, grant agreement No. 643476.

The VIBRE and COMBAT studies were supported by The Netherlands Organisation for Health Research and Development/The Netherlands Organisation for Scientific Research (ZonMw) under grants number 205100012 and 50-51700-98-120 respectively.

## Additional information

### Author contribution statement

S.M. and J.v.H. contributed equally to this work. S.M and JvH performed the systematic review, performed the experiments, interpreted the results and wrote the manuscript. N.W. contributed to bio-informatics analyses. M.A., D.M., J.P., M.d.J., the COMBAT consortium members, N.V., N.T.H. and C.S. contributed isolates and genome sequence data and associated meta data. M.d.J. and C.S. designed the study. All authors contributed to the writing of the manuscript.

